# AI-driven Classification of Cancer-Associated Fibroblasts Using Morphodynamic and Motile Features

**DOI:** 10.1101/2024.02.22.581611

**Authors:** Minwoo Kang, Chanhong Min, D. Somayadineshraj, Jennifer H. Shin

## Abstract

The heterogeneous natures of cancer-associated fibroblasts (CAFs) play critical roles in cancer progression, with some promoting tumor growth while others inhibit it. To utilize CAFs as a target for cancer treatment, issues with subtypes of CAFs must be resolved such that specific pro-tumorigenic subtypes can be suppressed or reprogrammed into anti-tumorigenic ones. Currently, single-cell RNA sequencing (scRNA-Seq) is a prevalent strategy for classifying CAFs, primarily based on their biomolecular features.

Alternatively, this study proposes assessing CAFs on a larger biophysical scale, focusing on cell morphological and motile features. Since these features are downstream effectors of differential gene expression combinations, they can serve as holistic descriptors for CAFs, offering a complementary strategy for classifying CAF subtypes. Here, we propose an artificial intelligence (AI) classification framework to comprehensively characterize CAF subtypes using morphodynamic and motile features. This framework extracts these features from label-free live-cell imaging data of CAFs employing advanced deep learning and machine learning algorithms.

The results of this study highlight the ability of morphodynamic and motile features to complement biomolecular features in accurately reflecting CAF subtype characteristics. In essence, our AI-based classification framework not only provides valuable insights into CAF biology but also introduces a novel approach for comprehensively describing and targeting heterogeneous CAF subtypes based on biophysical features.

## Introduction

The cancer-associated fibroblasts (CAFs) have risen to prominence as key players in cancer progression. These cells synthesize and remodel the ECM and also produce various growth factors. These functions modulate cancer metastasis and aid epithelial-mesenchymal transition (EMT). In addition, CAFs are involved in angiogenesis, modification of tumor metabolism, modulation of the immune system, and even drug access and therapy responses(1, 2). Accordingly, CAFs started to be recognized as reliable anti-cancer targets, so-called anti-CAF therapies.

CAFs were originally presumed to be homogeneous and accelerate tumor progression. However, it is now widely appreciated that CAFs are composed of heterogeneous populations, either pro-tumorigenic or anti-tumorigenic, and a growing body of evidence supports the multifaceted nature of CAFs(3, 4, 5, 6, 7, 8). Therefore, researchers in the field have been trying to identify subtypes of CAFs with molecular markers. For example, α-smooth muscle actin (*α-SMA*) is commonly used to identify CAFs. Other markers, such as fibroblast activation protein (*FAP*) and fibroblast-specific protein 1 (*FSP1*), also have been identified. However, those markers cannot exclusively demarcate the subpopulations and can be co-expressed in different subtypes. In order to utilize CAFs as a target for cancer treatment, understanding and classifying heterogenous CAF subtypes are critical such that specific pro-tumorigenic subtypes can be suppressed or reprogrammed into anti-tumorigenic ones(9).

Single-cell RNA Sequencing (scRNA-Seq) is a common practice to classify and study heterogeneous subpopulations of CAFs. However, as mentioned earlier, the so-called CAF markers are not exclusive, and the subtypes share molecular features, making it hard to identify subtypes solely from a molecular-level perspective(10, 11). To better delineate heterogeneous CAF populations, Bartoschek and colleagues introduced biophysical properties such as cell size, granularity, and scRNA-Seq data(11).

Similarly, Pelon and colleagues also used scRNA-Seq to classify subtypes of CAFs, and they identified four subtypes referred to as CAF-S1 to CAF-S4(4). Among the subtypes, the researchers focused on two subsets (CAF-S1 and CAF-S4) that exhibited a molecular characteristic of myofibroblasts, α-SMA. Despite sharing myofibroblastic features, the authors discovered notable functional differences between CAF-S1 and CAF-S4 on tumor progression, as well as significant differences in their cellular traction forces. Interestingly, although this aspect was not explicitly discussed in the paper, we are able to observe that these myofibroblastic CAF subsets displayed distinct morphological features. Specifically, CAF-S1 demonstrated a smaller cell spreading area with a larger aspect ratio, whereas CAF-S4 exhibited a larger spreading area with a smaller aspect ratio. This finding suggests that by describing CAF morphology in more detail, beyond simple geometric parameters like cell spreading area and aspect ratio, it may be possible to enhance the identification of heterogeneous CAF subtypes by incorporating morphological features in conjunction with biomolecular features.

CAFs’ motility, like their morphology, also reflects their heterogeneous nature. Costea and colleagues conducted a study on oral squamous cell carcinoma and identified two distinct CAF subtypes that exhibited different motile characteristics and varying effects on cancer cells(12). The CAF-N subtype, which displayed a transcriptomic profile similar to that of normal oral fibroblasts, had a higher proportion of migratory fibroblasts compared to the CAF-D subtype, which exhibited a more divergent transcriptomic profile compared to normal oral fibroblasts. These findings highlight the potential of incorporating morphological and motile features of CAFs alongside biomolecular features to provide a comprehensive description of CAF subtypes. The morphology and the motile characteristics of CAFs result from gene expression combinations. Thus, those characteristics can be holistic readouts to describe CAFs.

In addition, unlike biomolecular analysis, a fixed cell-based end-point assay, morphological or motile features of cells can be traced with live-cell imaging. That is, those features can provide information on the dynamic changes of CAFs. This approach, leveraging the temporal information of cells, has been demonstrated to be valuable in recent studies employing deep learning techniques(13, 14, 15, 16, 17).

In this study, we explore the utility of morphodynamic and motile features of CAFs as novel indicators of their heterogeneity, employing an AI-based classification framework. To achieve this, we propose an artificial intelligence (AI) classification framework specifically designed to classify CAF subtypes based on morphodynamic and motile features. Using deep learning multiple unsupervised and supervised machine learning algorithms, we extract the morphodynamic and motile features of cells from label-free live-cell imaging data of CAFs. Our approach, demonstrated using in vitro breast CAF models, co-cultured with two different breast cancer cell lines (luminal A, MCF-7 and triple negative, MDA-MB-231), reveals that these features effectively reflect the bimolecular characteristics of the CAF models.

## Results

### Validation of in vitro breast CAF models with CAF markers

To validate our hypothesis that morphodynamic and motile features of CAFs can be used as a complementary strategy to biomolecular features for a more comprehensive understanding of CAF heterogeneity, we developed and utilized in vitro breast CAF models as described in the materials and method section. Briefly, human dermal fibroblasts (HDFs) were co-cultured with breast cancer cell lines of different aggressiveness levels (Fig. 1a). Previous reports indicate that fibroblasts co-cultured with cancer cells in vitro were activated and demonstrated similar interactions with cancer cells as CAFs do(18, 19, 20). Another advantage of using in vitro CAF models is that we can obtain cells with similar activation patterns for multiple experiments, which provides consistency for analysis and is important for the verification of our new strategy to study CAF biology. To verify whether the in vitro breast CAF protocol induces the activation of fibroblasts, we investigated the expression levels of CAF markers in each fibroblast with qPCR (Fig. 1b). Except for *PDGFRβ* expression, the data revealed that the expression levels of +MCF-7 CAFs and +MDA-MB-231 CAFs were not statistically different from the control fibroblasts. It should be noted that PCR only provides the average expression level of individual cells in the samples, limiting the ability to elaborate on heterogeneous populations within the samples. However, even though the expression levels of *PDPN* in +MDA-MB-231 CAFs were not statistically higher than others, it is plausible that some fibroblasts in the +MDA-MB-231 CAFs were activated and expressed *PDPN* (*PDPN*^+^ CAFs). Likewise, we can infer that *FAP*^+^ CAFs were present in the two in vitro CAF models based on their higher expression compared to the control (*n.s: statistically non-significant.*). Furthermore, it is reasonable to say that the in vitro +MDA-MB-231 CAFs induced by the co-culture protocol reflected the in vivo breast CAFs well because *FAP*^+^*PDPN*^+^ CAFs (immunosuppressive phenotype) are found in dense ECM, which is the well-known characteristic of triple-negative breast cancer TME (21).

**Fig. 1.**
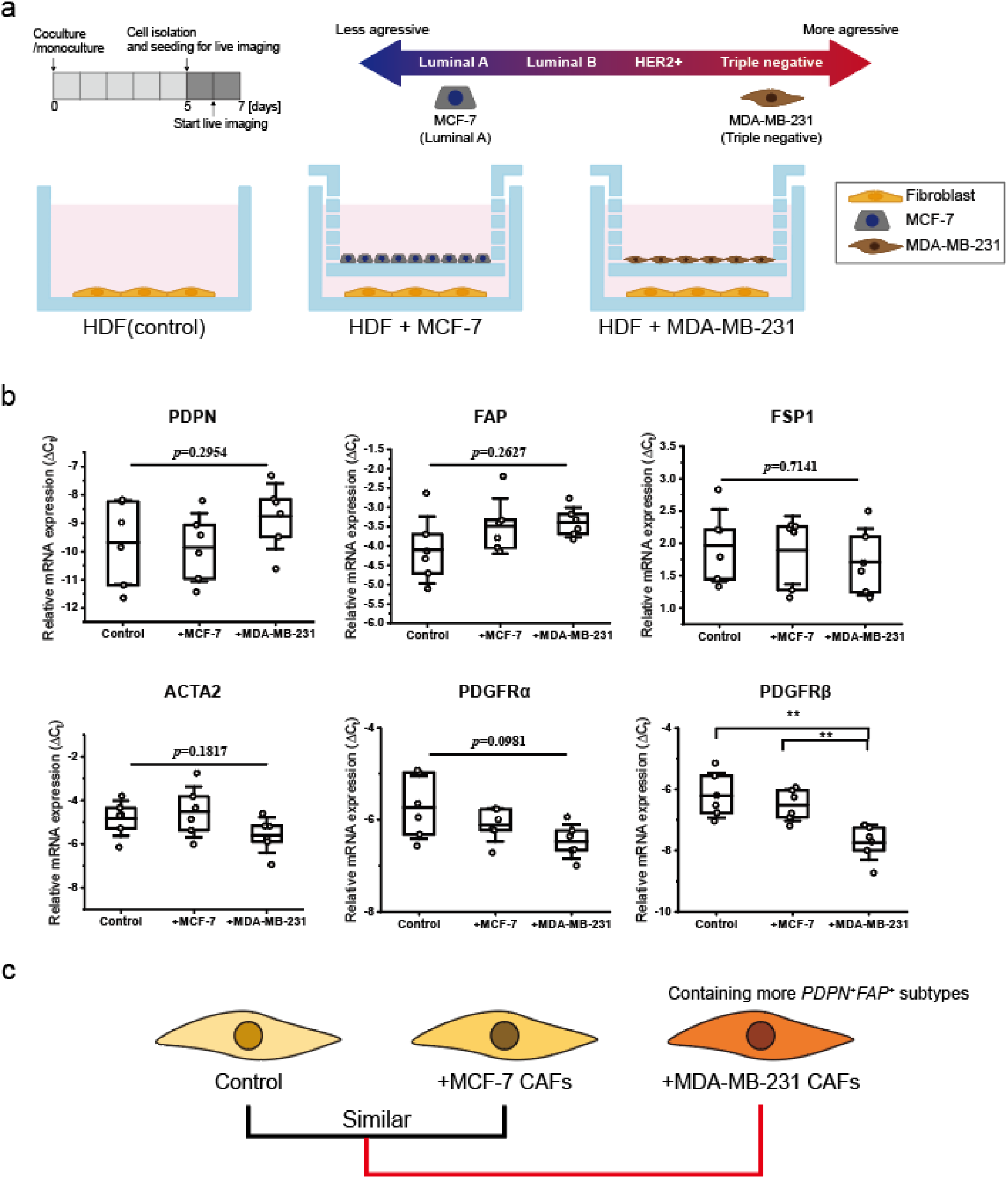
Establishing in vitro breast CAFs and validation with CAF markers. (a) In vitro breast CAFs were induced by co-culturing them with breast cancer cell lines of varying aggressiveness. (b) Relative mRNA expressions of CAF markers of normal fibroblasts and in vitro breast CAFs. (c) Assumed similarity between control and in vitro breast CAF models based on the biomolecular assay. Control, +MCF-7, +MDA-MB231 stand for normal fibroblasts, in vitro breast CAFs co-cultured with MCF-7, and in vitro breast CAFs co-cultured with MDA-MB-231 respectively. n = 6 and data represent mean ± SD.

On the other hand, the +MDA-MB-231 CAFs could be considered the most distinct group due to their deviated expressions compared to other groups (upregulated: *PDPN* (*n.s.*), downregulated: *ACTA2* (*n.s.*), *PDGFRα* (*n.s.*), *PDGFRβ (*\*\*), **: *p*<0.01). Conversely, the control group could be seen as the most heterogeneous in nature(22, 23). Therefore, considering the co-culture methods as inducing fibroblast activation in a specific direction, +MDA-MB-231 CAFs would be the most homogenous group containing *FAP*^+^*PDPN*^+^ CAFs. At the same time, +MCF-7 CAFs would more closely resemble the control (Fig. 1c). With this in mind, we characterized the heterogeneity of the cells through biophysical states, including morphodynamic and motile features.

### Morphodynamic features for CAF classification

Live imaging of single fibroblasts provides spatiotemporal information about cells. Originally, live imaging was conducted for 24 hours; however, we utilized image stacks from the initial 12 hours due to active proliferation after 12 hours of live imaging. We captured a total of ∼70,000 morphology states from four different live-cell imaging sessions (n=4). Supplementary Figure 1 shows the random single fibroblasts from normal fibroblasts and in vitro breast CAFs. From the figure, it was apparent that distinguishing each cell type solely by visual inspection was challenging. Therefore, we needed appropriate morphological descriptors to analyze the morphodynamic states of cells. Fundamentally, we calculated basic morphological variables of the cells from the live imaging dataset. In addition, the phase-contrast images of cells provide not only the geometry information but also texture information through light intensity differences. Also, Zernike moments, known for their orthogonal properties, noise robustness, and rotation and scale invariance, were utilized for their effectiveness in biomedical imaging applications(24, 25, 26, 27, 28, 29). To this end, we selected 52 morphological features from three morphological feature groups described in the materials and method section (basic morphology, texture features, and Zernike moments, Supplementary Figure 2) and extracted the features from every single fibroblast. While each individual morphological feature was capable of highlighting distinctions among cell groups (normal fibroblasts and in vitro breast CAFs, see supplementary figure 3), utilizing single features or dimensions is insufficient for comprehensively capturing the extensive diversity in cellular morphology. To overcome this, we integrated all 52 morphological features of each individual fibroblast, thus generating high-dimensional morphological features for each cell. To visualize high-dimensional morphological features, we performed dimensionality reduction with PCA. The first two principal components (PC1 and PC2) explained 22.6% and 16.3% of the variance, respectively, and the top 5 PCs and their most correlated variables were plotted (Fig. 2a). From the right panel of Fig. 2a, we observed that PC1 is highly associated with cell sizes (minor axis length), and PC2 is highly correlated with circularity, highlighting that the basic morphological features substantially influence the image representation of normal fibroblasts and in vitro breast CAFs compared to texture features and Zernike moments.

**Fig. 2.**
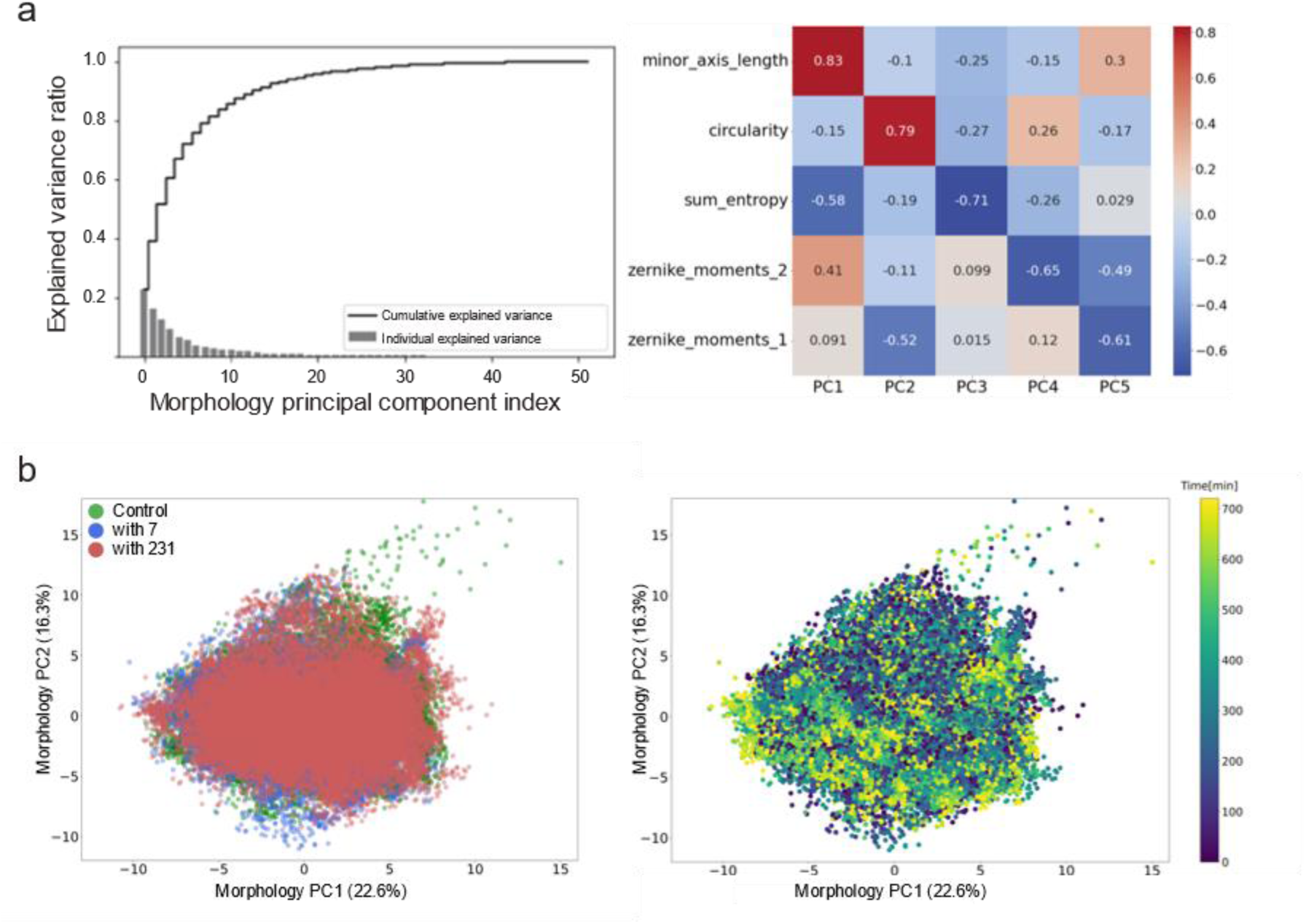
Embedding of morphology state of cells onto 2D morphology state space. (a) Dimensionality reduction of morphological features with principal component analysis (PCA). (Left panel) The individual variance is explained by principal components (PCs) and the cumulative explained variance. (Right panel) The top 5 PCs and their corresponding variables. (b) Embedding of morphology state of cells onto 2D morphology state space. (Left panel) Morphological states of normal fibroblast (control), +MCF-7 CAF (with7), and +MDA-MB-231 CAF (with231). (Right panel) Temporal fluctuation of morphological states.

The morphology states of all cells were then embedded into morphology PC1-PC2 state space (Fig. 2b). We observed that the morphological states of fibroblasts were very heterogenous, and each group (normal fibroblasts and in vitro CAFs) overlapped in most of the morphology state space, which was expected from CAF marker assay (Fig. 2b left panel). In addition, the morphology state showed temporal fluctuation, but identifying a pattern of temporal changes in morphology from the graph was challenging (Fig. 2b right panel). To quantitatively assess the heterogenous morphology states, we defined seven morphology states using k-means clustering (Fig. 3a). From the defined morphology states, we visualized the distribution of morphology states within the samples with a heatmap (Fig. 3b). We noticed that the control group and +MCF-7 CAF group showed similar distribution patterns from the heatmap, which was further supported by the unsupervised hierarchical clustering with Euclidean distance. Subsequently, the morphological heterogeneity of each group was represented by Shannon entropy (Fig. 3c). Since a lower value indicates a more homogeneous phenotype, the +MDA-MB-231 CAF, having the lowest Shannon entropy, appeared to be the most homogeneous group. Therefore, it was demonstrated that morphology features and the defined morphology states could effectively differentiate distinct CAF phenotypes.

**Fig. 3.**
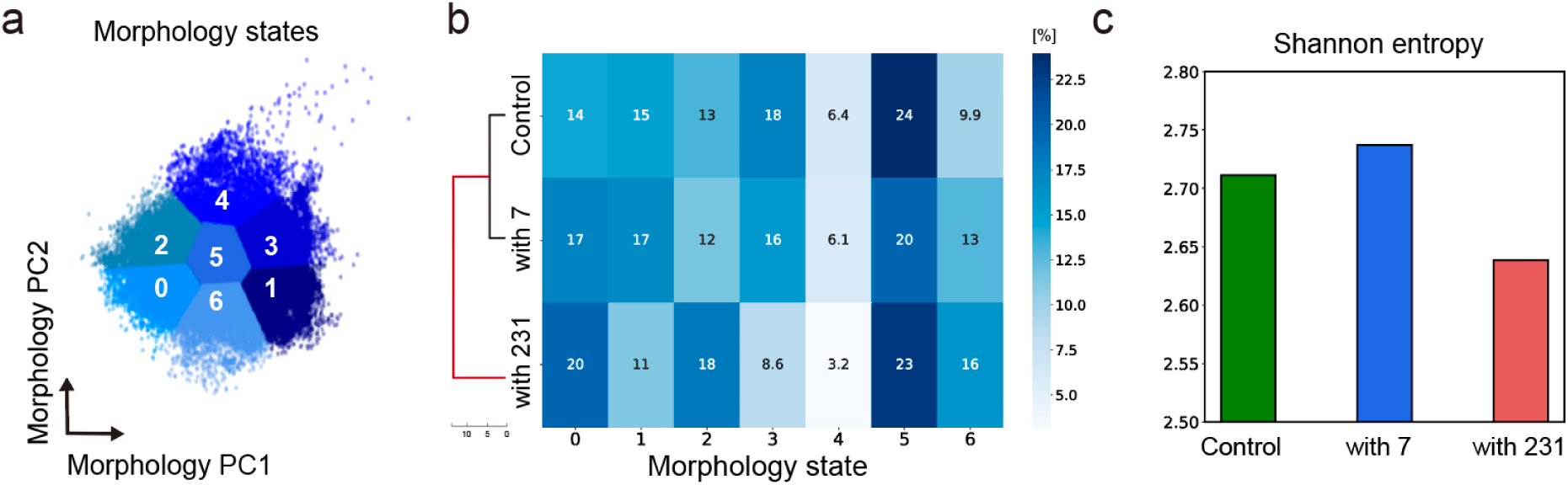
Defined morphology states in a 2D morphology state space and the distribution of morphology states of each group. (a) 7 defined morphology states identified by unsupervised clustering. (b) Defined morphology state distribution for each group and unsupervised hierarchical clustering of fibroblasts. (c) Morphological heterogeneity within the groups represented by Shannon entropy.

As mentioned earlier, we captured the temporal fluctuation of morphological features (Fig. 2c right). Therefore, we also visualized the distribution of morphology states of each group throughout the time axis (Fig. 4a). Even though the fluctuations of each state were visualized, gaining insight into the temporal variation of the morphology states remained challenging. To handle this problem, we tried to incorporate the temporal fluctuation of the morphology state, which led us to define morphodynamic states with time series k-means clustering. We defined three morphodynamic states (Fig. 4b). Similar to how we visualized the morphology states earlier, the distribution of morphodynamic states of fibroblasts was plotted with the heatmap (Fig. 4c). The morphodynamic states also revealed similar distribution patterns of control and +MCF-7 CAFs. This time, unsupervised hierarchical clustering with Euclidean distance further emphasized the greater distance between the +MDA-MB-231 CAFs and others, suggesting that the morphodynamic states more accurately reflect the biological features of fibroblasts and CAFs. However, the heterogeneity of +MDA-MB-231 CAFs and control was measured to be similar, failing to highlight the most homogenous groups (+MDA-MB-231 CAFs) containing the specific CAF types (*PDPN*^+^*FAP*^+^) (Fig. 4d). This result suggests that the morphologies captured at each time point are sufficient to classify the CAFs, and the changes of morphologies over time might not be as critical as the morphologies themselves in our experimental condition.

**Fig. 4.**
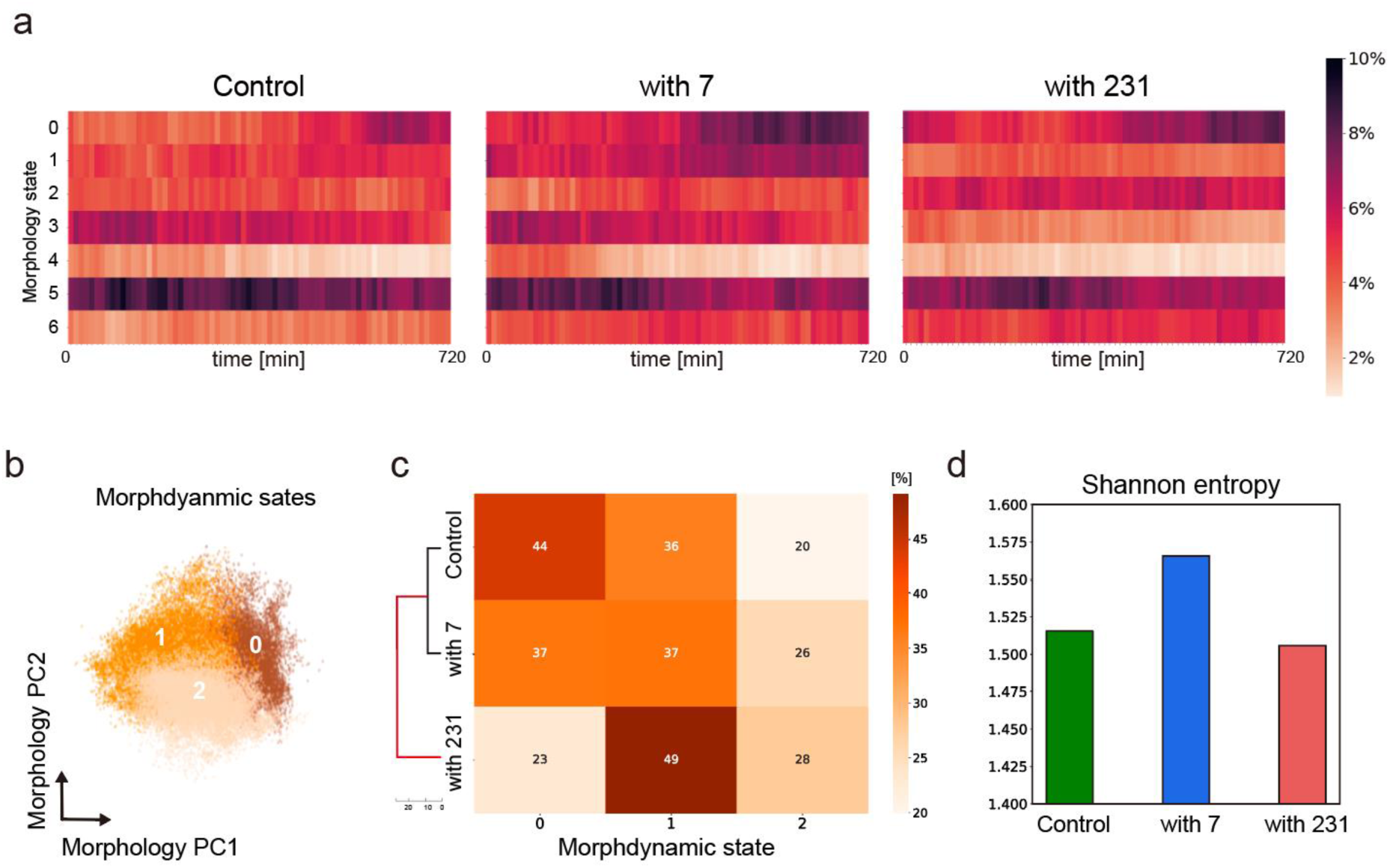
Defined morphodynamic states in a 2D morphology state space and the distribution of morphodynamic states of each group. (a) Temporal fluctuation of morphology states in each group. (b) 3 defined morphodynamic states by time series k-means clustering. (c) Defined morphodynamic state distribution of each group and unsupervised hierarchical clustering of fibroblasts. (d) Morphdynamical heterogeneity within the groups represented by Shannon entropy.

### Motile features for CAF classification

As we observed from the previous section, each individual cell within the group exhibited distinct morphology. Likewise, cells in a population also showed heterogeneous motile characteristics. In line with the approach of employing multiple features to quantify cellular morphology, we sought to comprehensively capture the diverse motile characteristics of fibroblasts. To achieve this, we specifically chose 50 motile features outlined in the materials and methods section, encompassing general features, MSD features, and random walk features. Subsequently, these features were extracted from each individual fibroblast (supplementary figure 4). Fibroblasts migrating in or out of the frame and proliferating ones during the live imaging were excluded from the analysis. Therefore, a total of ∼940 cells were analyzed. To visualize high-dimensional motile features, we implemented dimensionality reduction with PCA. The first two components (PC1 and PC2) explained 19.6% and 16.6% of the variance, and the top 5 PCs and the corresponding highest correlated variables were plotted (Fig. 5a).

**Fig. 5.**
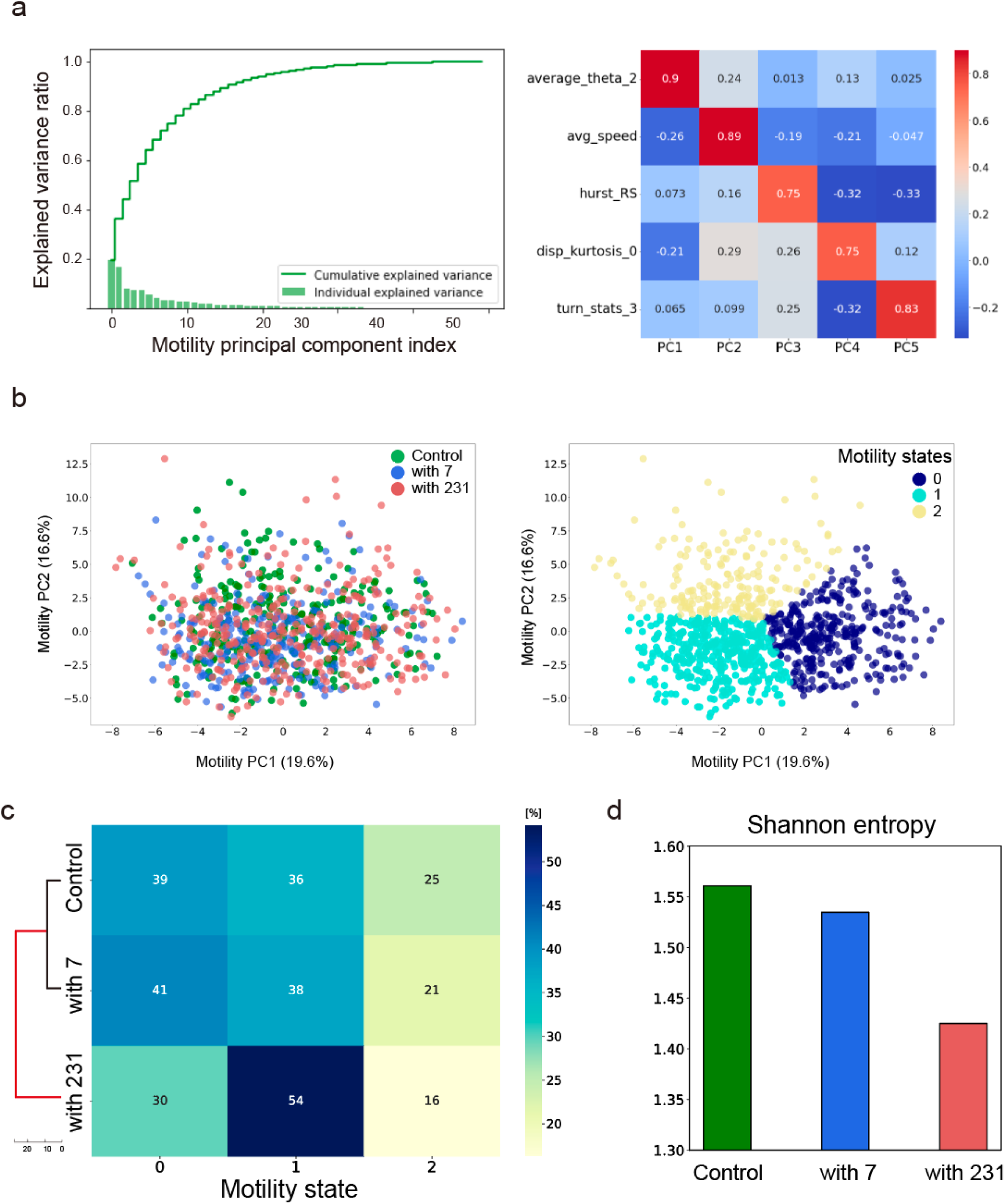
Embedding the motility state of cells into 2D motility state space, defining motility states, and the distribution of motility states of each group. (a) Dimensionality reduction of motility features with PCA. (Left panel) The individual variance explained by PCs, and the cumulative explained variance. (Right panel) The top 5 PCs and corresponding variables. (b) Embedding of motility state of cells onto 2D motility state space. (Left panel) Motility states of normal fibroblast (control), +MCF-7 CAF (with7), and +MDA-MB-231 CAF (with231). (Right panel) 3 defined motility states by unsupervised clustering. (c) Defined motility state distribution of each group and unsupervised hierarchical clustering of fibroblasts. (d) Motility heterogeneity within the groups represented by Shannon entropy.

From the right panel of Fig.5a, we observed a notable correlation where PC1 exhibited a strong association with the turning motion (average theta 2), and PC2 is correlated with speed. This observation implies that the general features among three motile feature groups (general features, MSD feature, and random walk features) contribute to the motile representation of normal fibroblasts and in vitro breast CAFs.

The motility states of all cells were then embedded in the motility PC1-PC2 state space (Fig. 5b left). We defined three motility states to quantitatively grasp the heteronomous spectrum of motility states by performing k-means clustering (Fig. 5b right). From the defined motility states, we quantified and visualized the distribution of fibroblasts with a heatmap (Fig. 5c). Clearly, the control group and +MCF-7 CAF group exhibited similar distribution patterns on the heatmap, which was corroborated by the unsupervised hierarchical clustering with Euclidean distance. The motile features also accurately identified the most homogenous group, +MDA-MB-231 CAFs, having the lowest Shannon entropy (Fig. 5d). Even though using only morphodynamic or motile features to describe the biophysical state of fibroblasts well classified the CAFs, it would be beneficial to incorporate the morphodynamic features into the motile features to define a more inclusive biophysical state of cells. In addition, embedding cell states onto PC1-PC2 space will not be enough to represent the data (the trade-off between information loss and dimensionality reduction). Therefore, we will combine morphodynamic and motile features in the next section to derive comprehensive morphodynamics-motility composite descriptors.

### Morphodynamics-motility composite descriptors for CAF classification

We established morphodynamics-motility composite descriptors by incorporating the morphodynamic features with motility features (Fig. 6a). To this end, we selected the top PCs that could explain 95% of each morphodynamic and motility features (top 20 morphology PCs and top 23 motility PCs). The morphology PCs were time-averaged to standardize the dimension. Then, the mean morphology PCs and motility PCs were combined as one feature vector. We used Uniform Manifold Approximation and Projection (UMAP) instead of PCA to visualize and embed the morphodynamics-motility composite features in two-dimensional space because the PCs from the composite feature vector exhibited a very subtle difference in explained variance (Supplementary Figure 5). Therefore, embedding the data into the first two components of PCA could not preserve the data information. In the two-dimensional UMAP space, which we defined as a morphodynamics-motility composite descriptor space, the composite descriptors of cells (UMAP coordinates of cells) were plotted by each group (Fig. 6b left panel). Subsequent k-means clustering defined three morphodynamics-motility composite descriptor states (Fig. 6b right panel). To verify whether UMAP preserved the morphodynamic and motility features appropriately, we plotted the cells labeled with morphodynamics-motility composite descriptor states onto the two-dimensional space of various features (mean morphology PC1, 2 and motility PC1, 2). Observing the figure, the data points appeared well-clustered by the composite descriptors in most feature spaces (Fig. 7a). We then proceeded to visualize the distribution of fibroblasts with a heatmap (Fig. 7b). The composite descriptors well captured the similarity between the control group and +MCF-7 CAFs. The overall heterogeneity of each group calculated by Shannon entropy also supported the result. The Shannon entropy of the control group and +MCF-7 CAFs showed similar values, and +MDA-MB-231 CAFs showed the lowest value, suggesting that the composite descriptors were successful in classifying the subtypes and identifying the most homogeneous group (Fig. 7c).

**Fig. 6.**
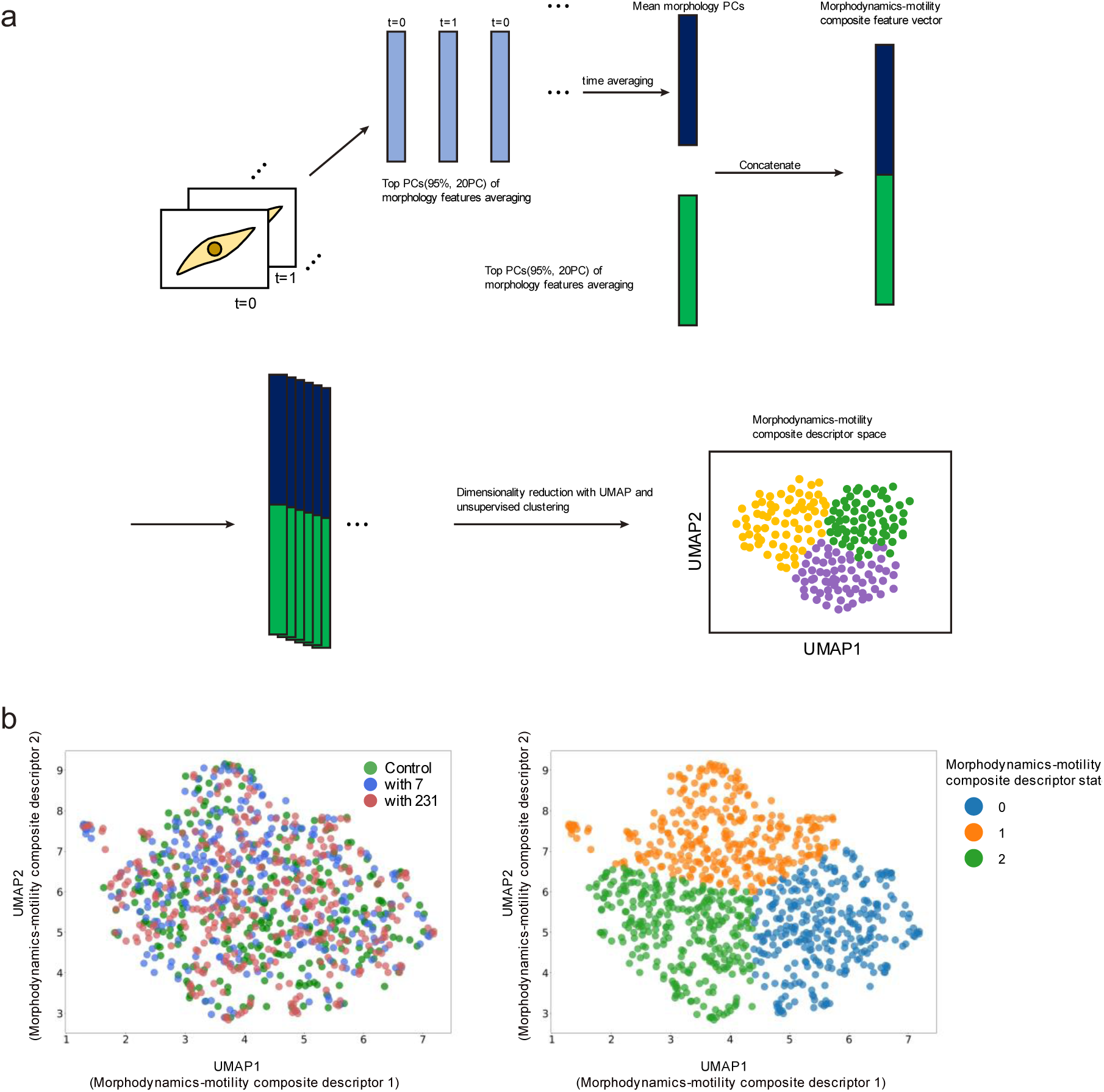
Establishing morphodynamics-motility composite descriptors and embedding the composite descriptors of cells onto the 2D composite descriptor space. (a) Establishing morphodynamics-motility composite descriptors by incorporating the morphodynamic features with motile features using machine learning algorithms. (b) Embedding of morphodynamics-motility composite descriptor state of cells onto a 2D morphodynamics-motility composite descriptor space. (Left panel) Morphodynamics-motility composite descriptors of normal fibroblast (control), +MCF-7 CAF (with7), and +MDA-MB-231 CAF (with231). (Right panel) 3 defined morphodynamics-motility composite descriptor states.

**Fig. 7.**
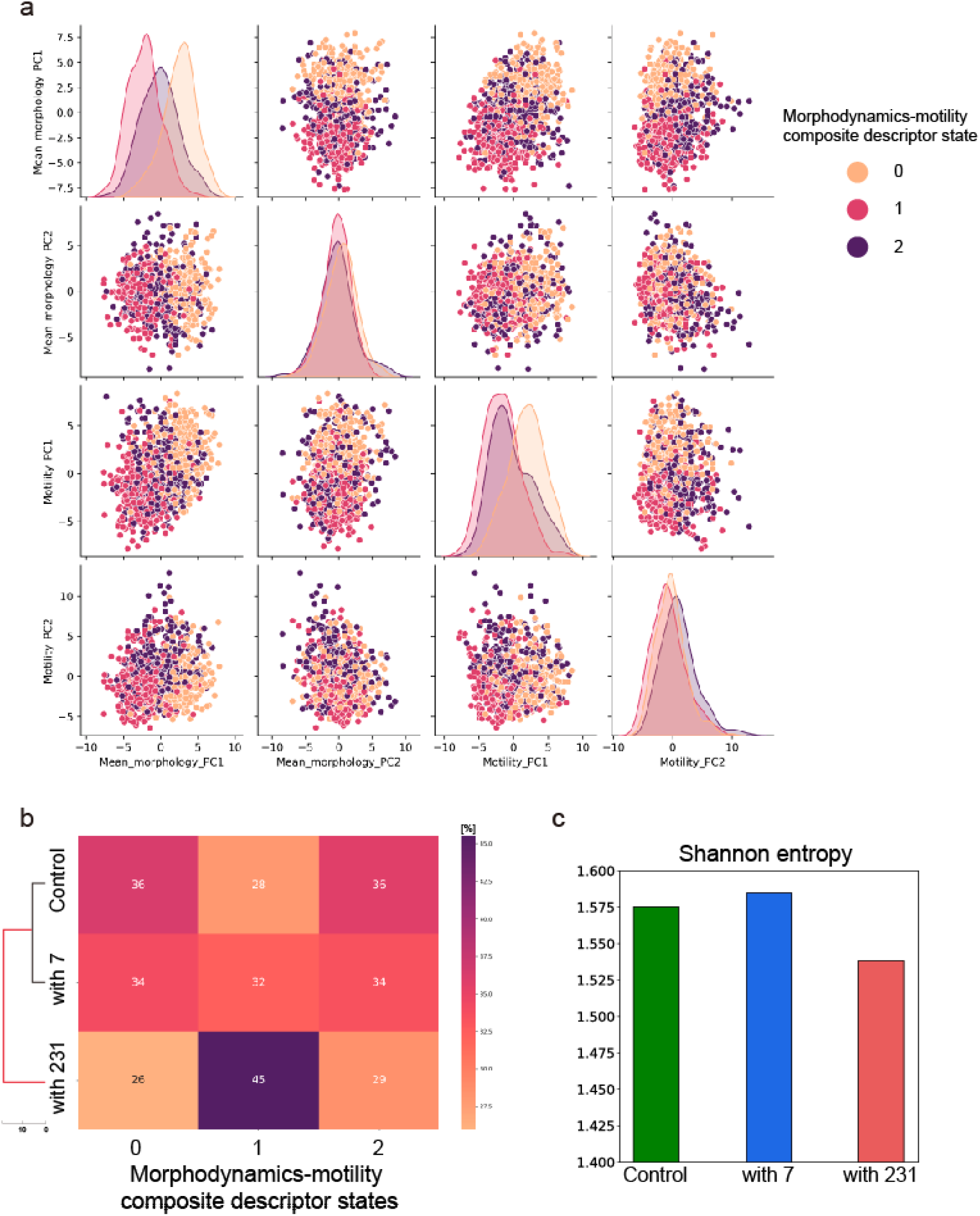
Verification of morphodynamics-motility composite descriptors and distribution of morphodynamics-motility composite descriptor states of each group. (a) Distributions of cells labeled with morphodynamics-motility composite descriptor states onto various feature spaces (mean morphology PC1, 2 and motility PC1, 2). The diagonal plots are kernel density estimation (KDE) of the univariate distribution of the corresponding features. (b) Defined composite descriptor states the distribution of each group and the unsupervised hierarchical clustering of fibroblasts. (c) Morphodynamics-motility heterogeneity within the groups represented by Shannon entropy.

## Discussion

CAFs exhibit heterogeneity that is fundamental to their role in the tumor microenvironment (TME). Traditional methods for identifying CAF subtypes have heavily relied on molecular biology approaches, particularly leveraging single-cell RNA sequencing (scRNA-seq) to delineate CAF subtypes. Nonetheless, our study pivots from the molecular to the biophysical realm, spotlighting morphodynamic and motile characteristics as key biophysical attributes reflective of underlying gene expression complexities. These biophysical features emerge as holistic indicators of cellular states, and their dynamic nature provides a more detailed view of cellular heterogeneity than static biomolecular assays.

In this context, we introduced a biophysical state-based AI Classification framework, with a special focus on in vitro breast CAFs as a test model. The use of in vitro breast CAFs is particularly advantageous as it allows for the generation of cells with consistent activation patterns across multiple experiments, lending reliability to the analysis crucial for validating our novel approach to studying CAF biology. Through co-culturing fibroblasts with breast cancer cell lines MCF-7 and MDA-MB-231, we established two distinct in vitro CAF models. Subsequent qPCR analysis for various CAF markers indicated a phenotype similarity between the control (normal fibroblasts) and +MCF-7 CAFs, while the +MDA-MB-231 CAFs exhibited a distinct activation profile of CAF markers, specifically with an upregulation of *PDPN* and *FAP*.

To perform a comprehensive biophysical state analysis for CAF classification, we integrated deep convolutional neural networks and other machine learning algorithms. Fifty-two morphological features were extracted from live-cell imaging and dimensionality reduction via PCA allowed us to visualize these features in a two-dimensional plane. Time-series k-means clustering identified three distinct morphodynamic states, encapsulating the temporal heterogeneity of cell morphologies. Heatmaps and unsupervised hierarchical clustering analyzed the distribution of these morphodynamic states, and Shannon entropy was employed to evaluate morphodynamical heterogeneity. Although the morphodynamic features could depict the similarity between the control group and +MCF-7 CAFs, it failed to predict the most homogeneous +MDA-MB-231 CAFs.

Further, we extracted fifty motile features and subjected them to similar dimensionality reduction and clustering analyses. These machine learning approaches enabled us to define three motility states and, as anticipated, the cells labeled with motility states were accurately classified into subtypes by unsupervised hierarchical clustering. To synthesize the morphodynamic and motile biophysical states, we combined these features into morphodynamics-motility composite descriptors for a more comprehensive CAF classification. These composite descriptors not only preserved the distinct morphodynamic and motile features but also effectively differentiated the biologically diverse CAF groups, pinpointing the +MDA-MB-231 CAFs as the most homogeneous subset (Fig. 8).

**Fig. 8.**
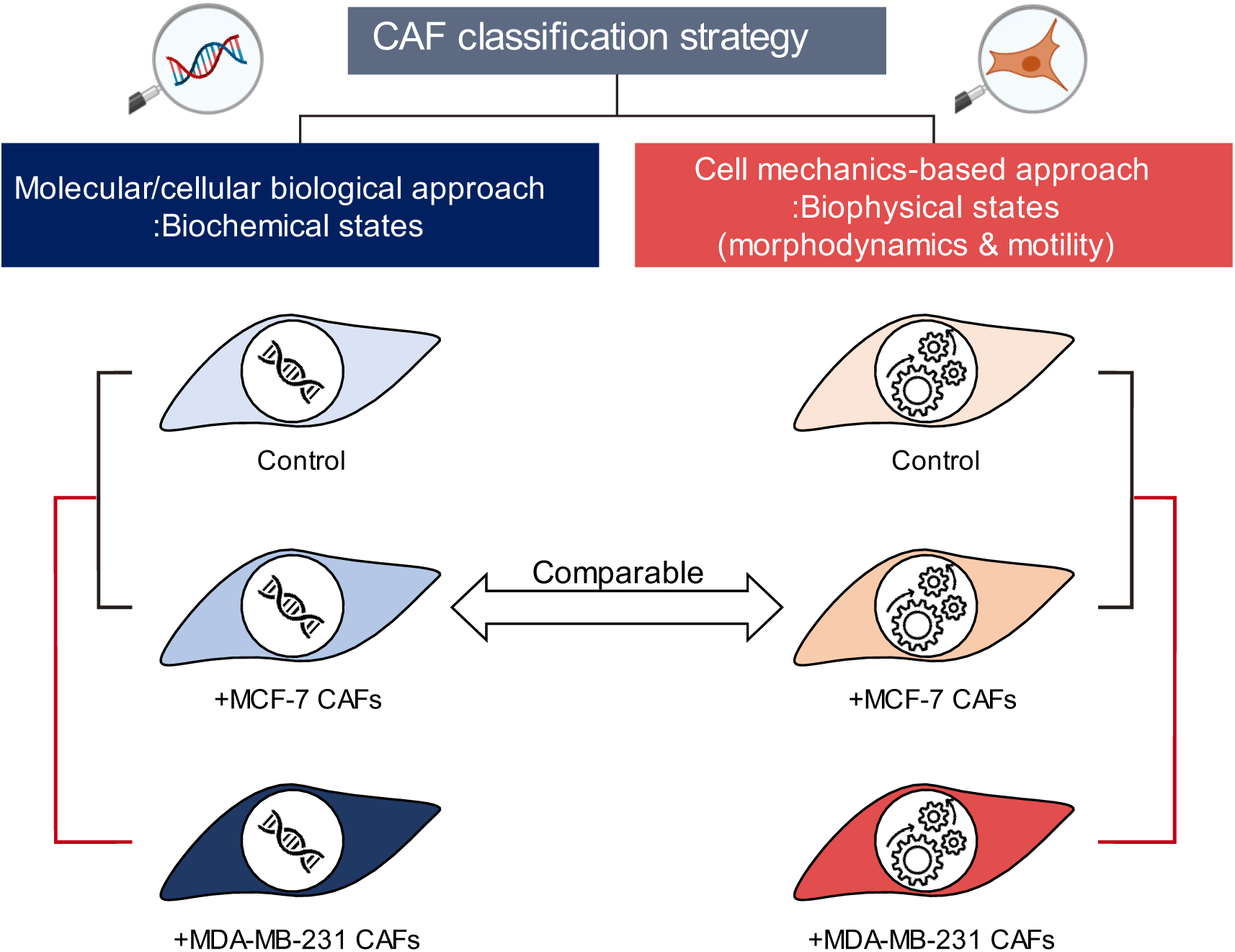
Comparison between different CAF classification strategies. Biophysical state-based CAF classification predicted the same result as biochemical state-based classification.

Our findings offer a novel perspective in CAF research, demonstrating a state-of-the-art biophysical state-based AI framework that capitalizes on morphodynamic and motile features for the meticulous classification of CAFs. This approach could potentially revolutionize our understanding and treatment strategies for cancer. Nonetheless, it is crucial to acknowledge the limitations of our study. Our verification of biomolecular signatures in in vitro CAFs was confined to qPCR analysis, providing only a bulk expression profile of the cells. A comparative analysis incorporating these biophysical states (composite descriptors) alongside transcriptomic data (scRNA-seq) is essential to further authenticate and potentially augment the relevance of the morphodynamics-motility composite descriptors in the realm of personalized medicine.

## Materials and Methods

### Cell culture and establishing in vitro breast CAF models

Human adult dermal fibroblasts (HDFs, Lonza), MCF-7, and MDA-MB-231cells were cultured separately and maintained in DMEM (WelGene) supplemented with 10% fetal bovine serum (FBS, WelGene), and 1% penicillin-streptomycin (WelGene). To make different in vitro breast CAFs, HDFs were co-cultured with either MCF-7 or MDA-MB-231 in a co-culture dish (SPL, 209260) for five days. The seeding density for HDFs was 4,650 cells/cm^2^, and cancer cells were 2,325 cells/cm^2^ (Fig. 1a). To verify whether the short days (5 days) of co-culture induced the CAF phenotypes, real-time qPCR was performed with several CAF markers.

### Live-cell imaging of single fibroblasts

Live-cell imaging was performed on the Axio Observer.Z1/7 (Carl Zeiss) microscope equipped with a climate-controlled chamber (37 °C and 5% CO2). Phase-contrast images of each single-fibroblasts were taken every 10 min for 24 hours using a 5x objective lens and 1x optovar magnification changer. Fibroblasts from each condition (monoculture or co-culture with cancer cells) were seeded with the sparse seeding density of 2,000 cells/well on 6-well plates to capture singe-cells and minimize overlapping cells while migrating

### Instance segmentation, tracking, and extracting morphodynamic and motile features of single cells

For the instance segmentation of single fibroblasts, we first performed semantic segmentation using U-net with the adapted backbone(30). To make training data for deep learning, we manually annotated the fibroblast in the images. A total of 26 random images containing 1,016 fibroblasts were annotated. The training data, which has a size of 1792×1024 pixels, were cropped into small patches (256×256 pixels). The total 728 patches of phase-contrast images and the corresponding annotated images were then randomly flipped, rotated, transposed, distorted, brightness changed, and contrast adjusted as a data augmentation technique for efficient learning. Then, the augmented 50,000 patches were used as training input. Resnet34 was used as the encoder block of the adapted U-net (Supplementary Figure 6)(31). The deep convolutional neural network (DCNN) was trained with the combination loss (binary cross entropy + Jaccard loss) and Adam optimizer. The parameters for the training were batch size 32 and epochs 100. The criterion to evaluate the learning was the IoU (intersect over union) score, and the IoU score greater than 0.5 is generally considered good. At the end of the training, the validation IoU score reached over 0.7 (Supplementary Figure 7a). However, the training IOU was more significant than the validation IOU, suggesting the overfitting of the learning. This might occur because live imaging intrinsically cannot capture the full spectrum of all possible cellular geometries due to its inherent limitations. Therefore, we evaluated the segmented images visually to check whether the overfitting caused the problem predicting cells. To this end, the small patches (256×256 pixels) were integrated to make the original image size (1792×1024 pixels). Even though there were some mispredicted areas, we confirmed that the model was trained well (Supplementary Figure 7b).

Binary cell images obtained from the semantic segmentation were then manually inspected and corrected, while the cells smaller than 200 pixels (not a cell) or touching the boundary of the image frame (partial cellular images) were deleted automatically in the process of instance segmentation. Then, we put bounding boxes for each cell to label cells in the frame. At the same time, 52 morphological features (basic morphology, texture features, and Zernike moments) of the labeled cells in each frame were extracted (Supplementary Figure 2). The basic morphology features consisting of 10 variables (area, extent, perimeter, minor axis length, major axis length, equivalent diameter, solidity, aspect ratio, circularity, compactness) were obtained using the scikit-image library(32). The texture features and Zernike moments of the cells were calculated using the Mahotas library(33).

To track the single cells of the image stacks, the previously labeled cells in each frame had to be marked with the same labeling all across the frames. The first step is matching cells with two adjacent frames, and this problem can be a linear assignment problem (LAP). Because we obtained the centroids and sizes of labeled cells from the previous instance segmentation, the matching problem can be done with a pairwise cost matrix, which is calculated by the centroid displacements and cell size difference (expressions 1 and 2). Therefore, one can see that the lowest cost means the proper matching of cells of adjacent frames. We set threshold values for distance (100 pixels) and size (2 folds) as the previous report did(34). The next step is relabeling the cells of all frames. In this step, there are three possible scenarios. First, if there is a match between two adjacent frames, we change the labels in the second frame with the labels of the first frame. Second, if the labels in the first frame are not found in the second frame, that is the end of the label. Lastly, if a new object exists in the second frame, we impose new labels (number). Therefore, if we relabel the cells from the first frame of the entire image stack, we sequentially relabel the adjacent frame by the final frame of the image stack and eventually correctly label all the cells in the image sequence (Supplementary Video 1).

To extract the motile features, centroids of cells in each time frame were extracted. Then, the 50 motility features were calculated by incorporating the Heteromotility tool (https://github.com/cellgeometry/heteromotility(35)) into our lab-made code. The motile features can be classified into three groups based on the class in the Python code of Heteromotility tools, which are general, mean square displacement (MSD), and random walk features (Supplementary Figure 4). The 33 general features were extracted in the study (net distance, linearity, Spearman’s rho2, progressivity, min speed, max speed, avg speed, time spent moving 0∼2, turn stats 0∼4, avg moving speed 0∼2, number of turns (average theta 0∼4), magnitude of turns (min theta 0∼4, max theta 0∼4)). From MSD feature, α values were extracted, which can be random motion, directed motion, or confined motion if the values are one, greater than one, or smaller than one, respectively. Finally, from the random walk features, 16 features were extracted as follows: difference in linearity, difference in net distance, displacement kurtosis, displacement skewness, displacement variance, Hurst exponent RS, Non-Gaussian coefficient (alpha2), autocorrelation 1∼4, partial autocorrelation 0∼4.

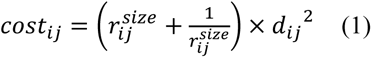

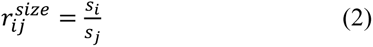

*d_ij_*∶ distance between the centroid of cell *i* and cell *j* from two adjacent frames

*s_i_* ∶ the size of cell *i*

### Dimensionality reduction of morphological and motile features with principal component analysis (PCA)

After extracting 52 morphological features of cells, PCA was performed. A total of 70,018 morphology states were embedded in two-dimensional morphology state space (morphology PC1-PC2). The same procedure was applied to the motile features of cells, and a total of 937 cells’ motile features were embedded in the two-dimensional motility state space.

### Unsupervised clustering to define biophysical states

To identify the heterogenous subtypes of fibroblasts in the two-dimensional biophysical state space, an unsupervised clustering algorithm, k-means clustering, was implemented. To obtain the optimum number of clusters, the elbow method or silhouette coefficient was used. For the clustering of time-series data, time-series k-means clustering with dynamic time warping (DTW) was implemented.

After defining the clusters (the defined biophysical states), the fibroblasts’ distribution of clusters was calculated and visualized with a heatmap. Then, unsupervised hierarchical clustering with average-linkage and Euclidean distance was carried out. Shannon entropy was used to quantify the general heterogeneity of fibroblasts (expression 3).

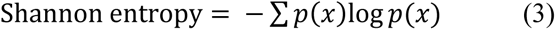

Where *p(x)* is the % distribution of the clusters

### Reverse transcription Quantitative PCR

After co-culturing for five days, the total RNA of fibroblasts was isolated with RNAiso reagent (Takara Bio, Japan) according to the manufacturer’s instructions. Extracted RNAs were reverse transcribed to cDNA using iScript cDNA Synthesis Kits (Bio-Rad, USA) and Biometra T-personal Thermal Cycler for the synthesis. Real-time qPCR was carried out in duplicates with iQ SYBR green supermix (Bio-Rad, USA) and a Bio-Rad CFX96 real-time detection system. Glyceraldehyde 3-phosphate dehydrogenase (GAPDH) was used for the reference gene. ΔCt values were used for hypothesis testing used to express relative mRNA expression normalized to GAPDH. ΔCt = Ct(reference gene) - Ct(gene of interest), Ct: Threshold cycle.

The following primers were used:

GAPDH (For: CTGGGCTACACTGAGCACC, Rev: AAGTGGTCGTTGAGGGCAATG) ACTA2 (For: CTATGAGGGCTATGCCTTGCC, Rev: GCTCAGCAGTAGTAACGAAGGA), FAP (For: TGAACGAGTATGTTTGCAGTGG, Rev: GGTCTTTGGACAATCCCATGT), FSP1 (For: GATGAGCAACTTGGACAGCAA, Rev: CTGGGCTGCTTATCTGGGAAG), PDPN (For: AACCAGCGAAGACCGCTATAA, Rev: CGAATGCCTGTTACACTGTTGA), PDGFRα (For: TTGAAGGCAGGCACATTTACA, Rev: GCGACAAGGTATAATGGCAGAAT), PDGFRβ (For: TGATGCCGAGGAACTATTCATCT, Rev: TTTCTTCTCGTGCAGTGTCAC),

### Statistical analysis

One-way ANOVA with Games-Howell post hoc tests was used for statistical hypothesis testing. P-values < 0.05 were considered statistically significant. Statistical tests were performed using *jamovi* version 1.6.23.0 (The jamovi project (2021), https://www.jamovi.org).

## Acknowledgment

We express our profound gratitude to Sang Yoon Oh, Tae Yoon Kwon, Hyuntae Jeong, and Myung Chul Kim for their invaluable contributions and productive discussions in shaping the development of the code. This research was supported by National Research Foundation granted by the Korean Government (NRF-2021R1A2C3008408).

## Author contributions

M.K. conceived the study and designed experiments. M.K. and C.M developed the code for AI classification. M.K., S.D. performed the experiments. M.K., S.D. and J.S. interpreted data.

M.K. wrote the draft. All authors edited the manuscript.

## Data availability statement

Python code for instance segmentation, tracking, and extracting morphodynamic and motile features of single cells is available at https://github.com/shinlab-kaist/CAF-project

