## Supplementary figures for "AI-driven Classification of Cancer-Associated Fibroblasts Using Morphodynamic and Motile Features"

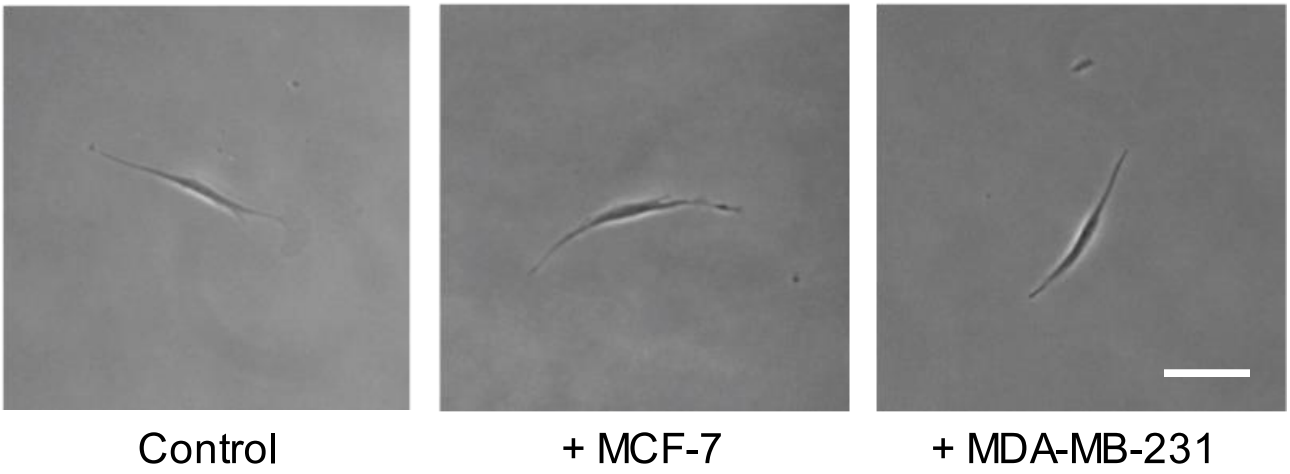


**Supplementary Figure 1 Random snapshots of normal fibroblasts and in vitro breast CAFs from live imaging.** Control, +MCF-7, and +MDA-MB-231 stand for normal fibroblasts, in vitro breast CAFs cocultured with MCF-7, and in vitro breast CAFs cocultured with MDA-MB-231, respectively. Scalebar: 100 µm.

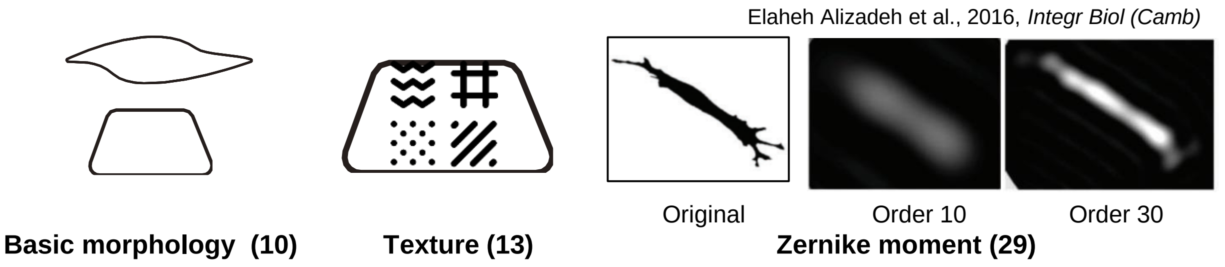


**Supplementary Figure 2 Total of 52 morphological features consisting of basic morphology, texture, and Zernike moment features.**

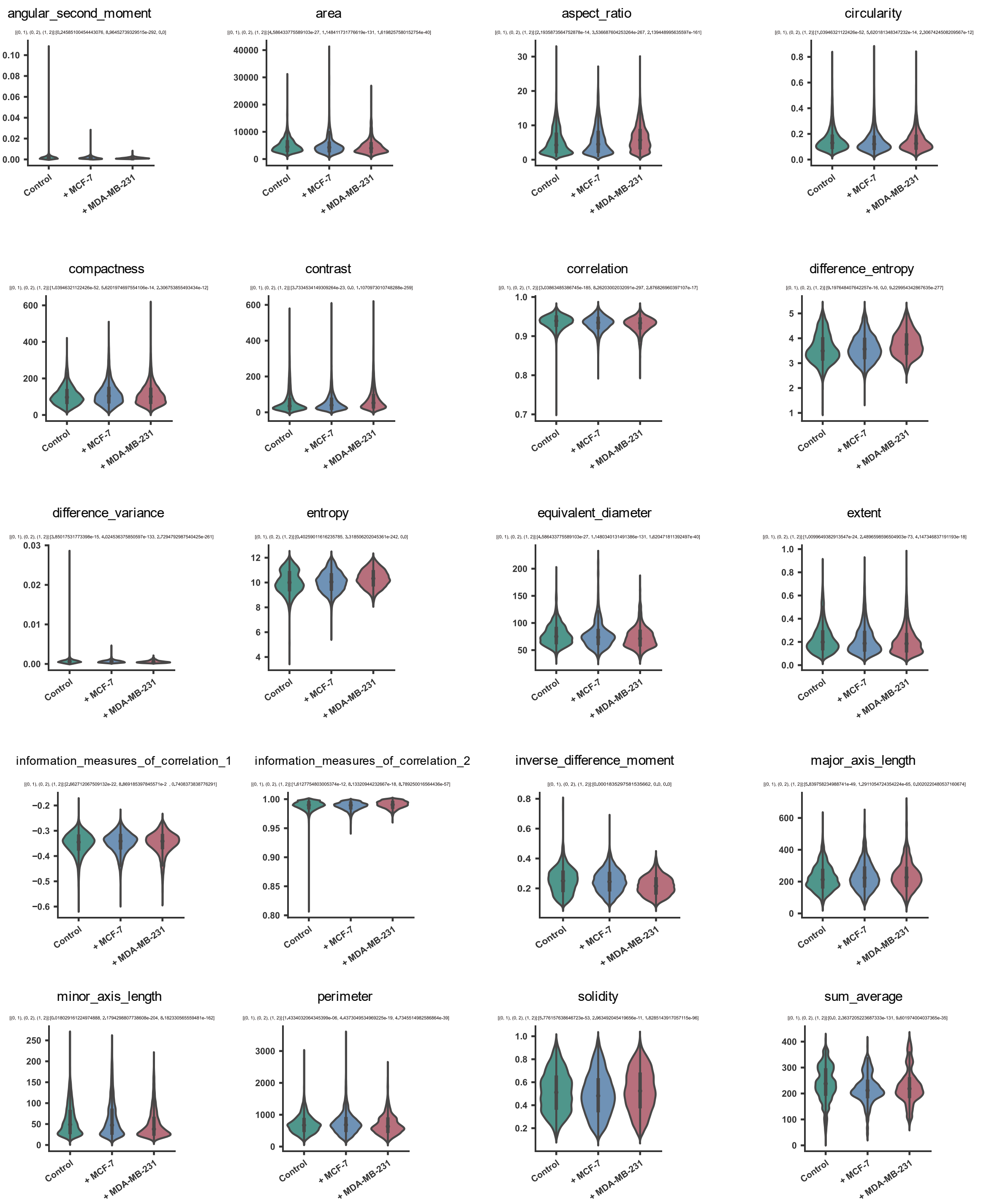
angular_second_moment
area
aspect_ratio
circularity
compactness
contrast
correlation
difference_entropy
difference_variance
entropy
equivalent_diameter
extent
information_measures_of_correlation_1
information_measures_of_correlation_2
inverse_difference_moment
major_axis_length
minor_axis_length
perimeter
solidity
sum_average
Control
+ MCF-7
+ MDA-MB-231

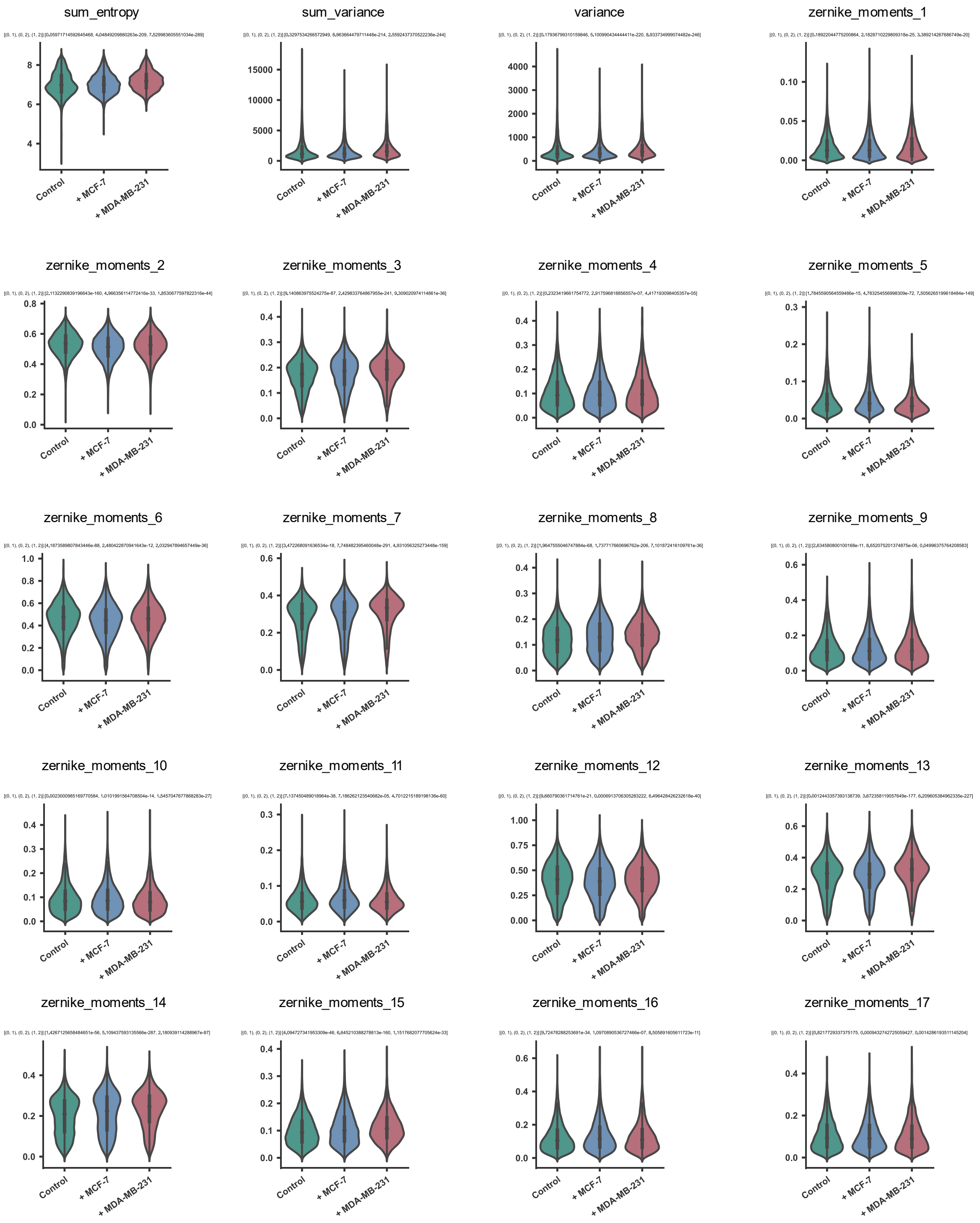

sum_entropy
sum_variance
variance
zernike_moments_1
zernike_moments_2
zernike_moments_3
zernike_moments_4
zernike_moments_5
zernike_moments_6
zernike_moments_7
zernike_moments_8
zernike_moments_9
zernike_moments_10
zernike_moments_11
zernike_moments_12
zernike_moments_13
zernike_moments_14
zernike_moments_15
zernike_moments_16
zernike_moments_17
Control
+ MCF-7
+ MDA-MB-231

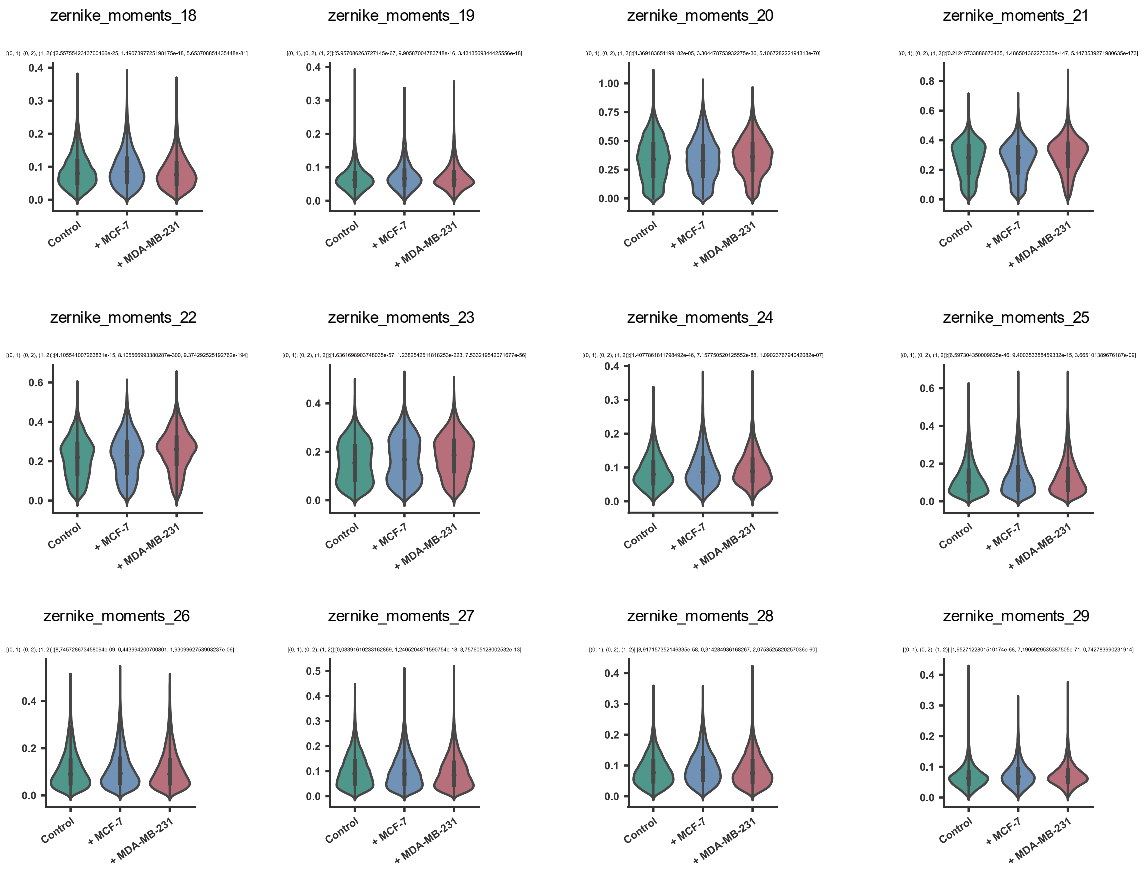


**Supplementary Figure 3  Comparative morphological analysis of normal fibroblasts and in vitro breast CAFs.** Figures illustrate individual morphological features among normal fibroblasts (Control), fibroblasts co-cultured with MCF-7 breast cancer cells (+MCF-7), and fibroblasts co-cultured with MDA-MB-231 breast cancer cells (+MDA-MB-231). The pairwise comparisons, denoted by the numerical pairs (e.g., (0,1)), assess differences between the Control group and the +MCF-7 CAFs. The values within the brackets represent the p-values derived from the Mann-Whitney U test, indicating the level of statistical significance for each comparison.

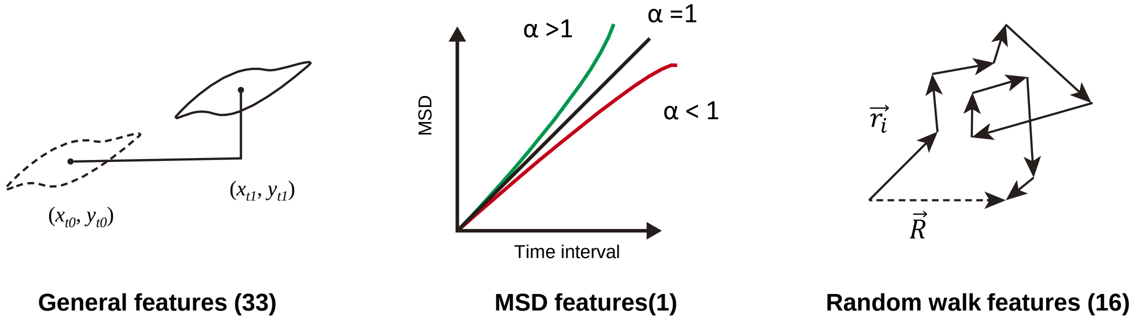


**Supplementary Figure 4  Total of 50 motile features consisting of general, MSD, and random walk features.**

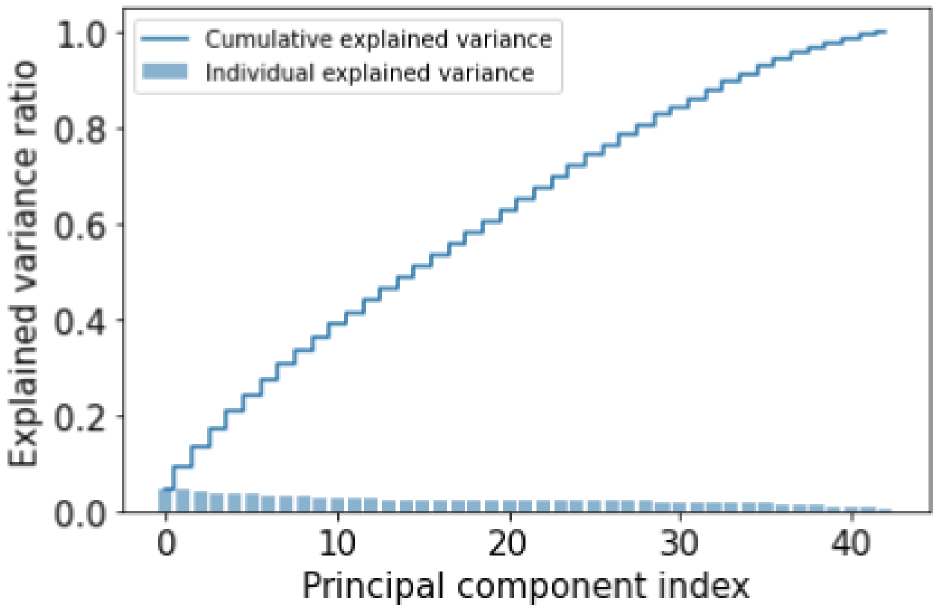


**Supplementary Figure 5 Establishing morphodynamics-motility composite descriptors and embedding of the composite descriptors of cells onto the 2D composite descriptor space.** (a) Establishing morphodynamics-motility composite descriptors by incorporating the morphodynamic features with motile features using machine learning algorithms.

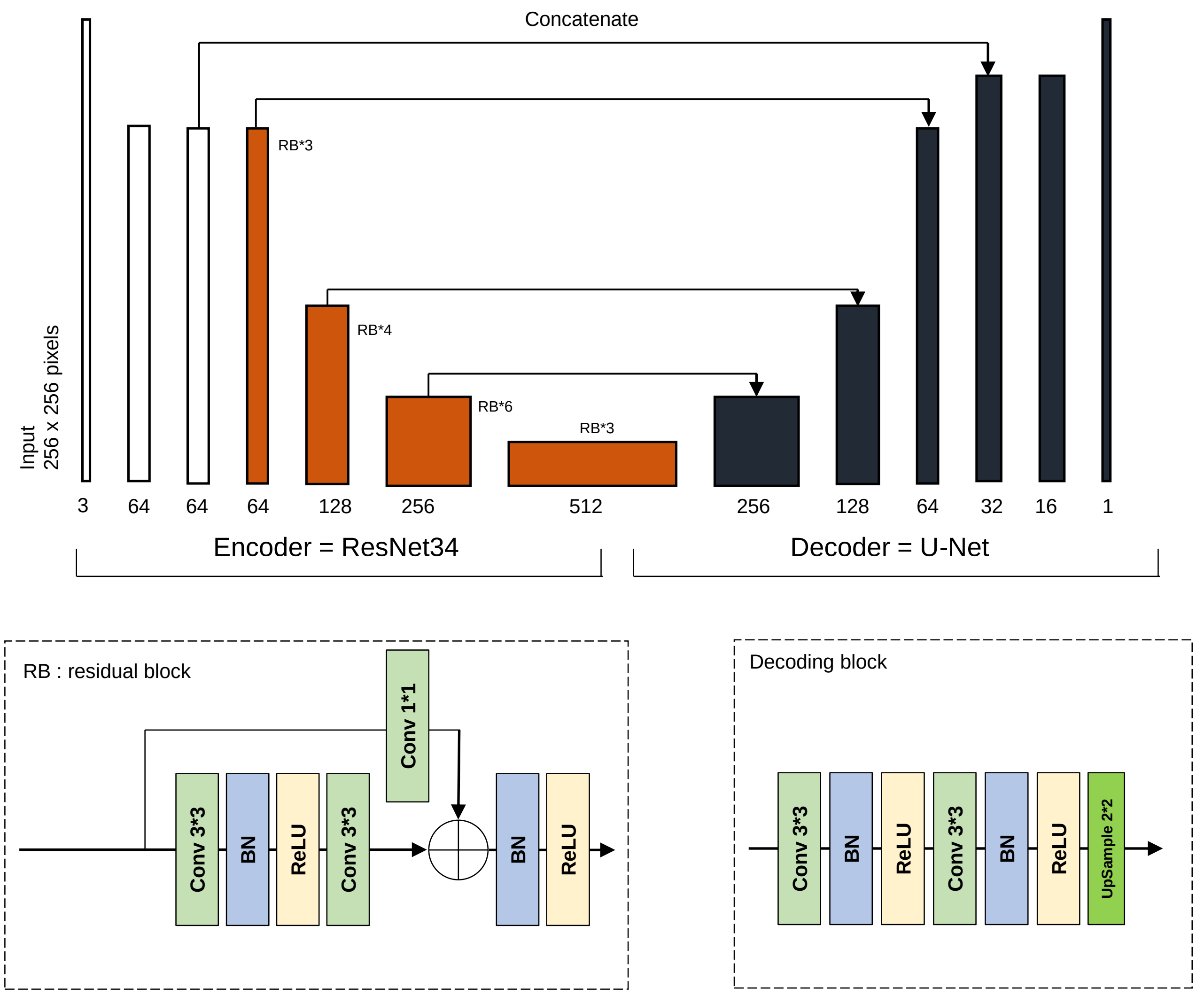


**Supplementary Figure 6 Architecture of the deep convolutional neural network for semantic segmentation of single cells.** Conv: convolution, BN: batch normalization, ReLU: Rectified linear unit.

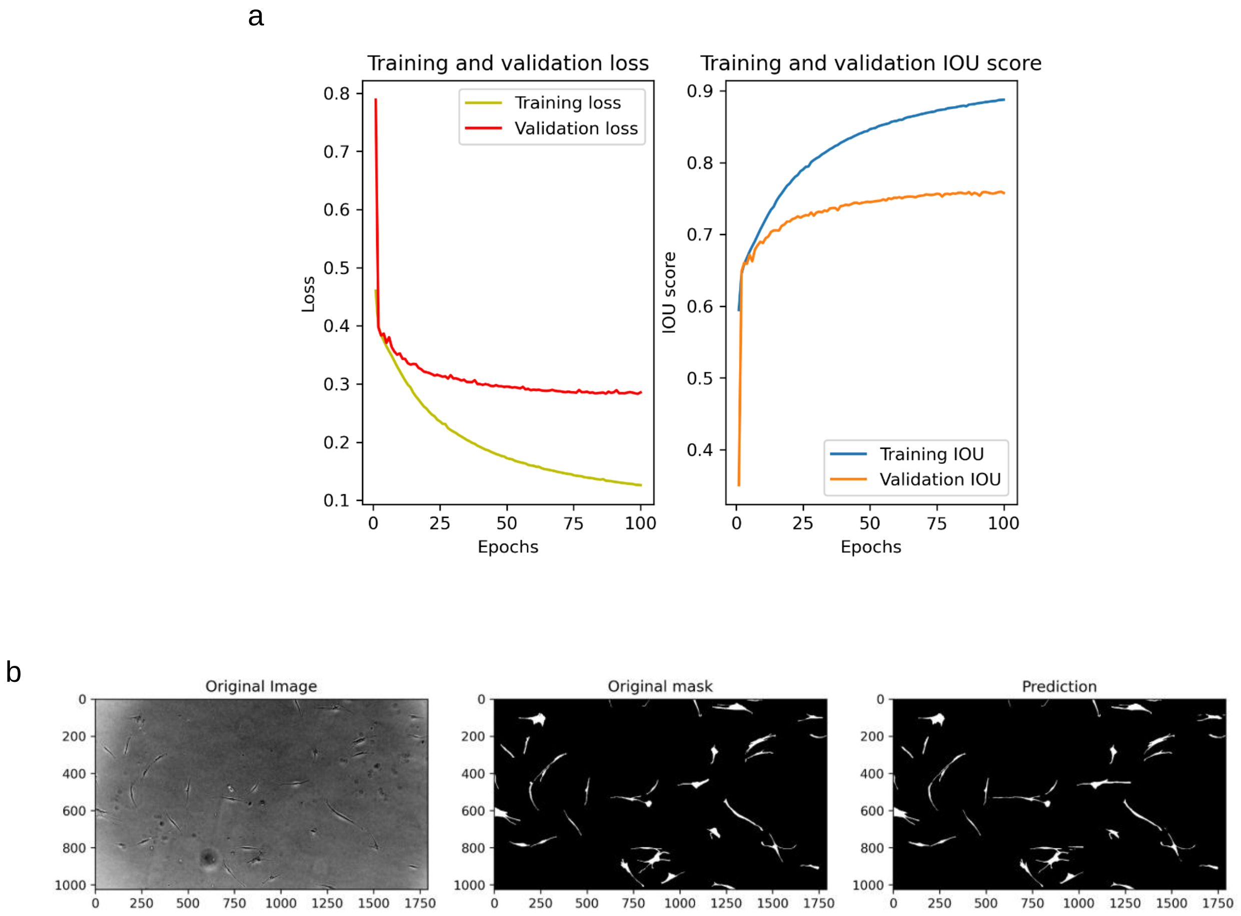


**Supplementary Figure 7 Validation of the training.** (a) Loss curve and IOU score of training and validation data set. (b) Visual comparison between the annotated image and the predicted image of fibroblasts.
